# REGENERATIVE GENE THERAPY IN THE HYPOTHALAMUS PROLONGS FERTILITY IN FEMALE RATS

**DOI:** 10.1101/2023.05.24.542027

**Authors:** Maria D. Gallardo, Mauricio Girard, Araceli Bigres, Marianne Lehmann, Silvia S. Rodriguez, Rodolfo G. Goya

## Abstract

There is substantial evidence that age-related ovarian failure in rats is preceded by abnormal responsiveness of the neuroendocrine axis to estrogen positive feedback. In middle-aged (M-A) female rats, we have demonstrated that intrahypothalamic gene therapy for insulin-like growth factor-I (IGF-I) started at 6 months of age extends the regular cyclicity of the animals beyond 10 month (the age at which MA rats stop ovulating) and preserves the integrity of the ovarian structure. Here, we implemented long-term regenerative gene therapy in the hypothalamus of young females. The goal was to extend fertility in the treated animals. We constructed a helper-dependent adenovector that harbors the green fluorescent protein (GFP) reporter gene as well as a gene tandem, termed STEMCCA, which harbors the 4 Yamanaka genes (oct4, sox2, klf4, and c-myc, OSKM), both under the control of a Tet-Off bidirectional promoter. An adenovector that only carries the gene for GFP was used as control. At 4 months of age 12 female rats received an intrahypothalamic injection of our OSKM-GFP vector (treated rats); 12 control rats a vector expressing GFP only (control rats). At 9.3 months of age control and treated rats were mated with young males. A group of 12 young intact female rats was also mated. The rate of pregnancy recorded was 83%, 8.3% and 25% for young, M-A control and M-A treated animals, respectively. Average litter size was 9, 3 and 3 for the corresponding groups. Mean pup BW was slightly higher in the M-A rats. In a preliminary study we confirmed that ovulation in our M-A females ceases at 10 months of age. Our results are in line with the evidence that viral vector-mediated delivery of the Yamanaka genes in the brain has strong regenerative effects without adverse side effects. The particular significance of the present results is that, for the first time, they show that long-term OSKM gene therapy in the hypothalamus is able to extend the functionality of such a complex system as the hypothalamo-pituitary-ovarian axis.

## INTRODUCTION

### Reproductive aging

In middle-aged female (MA) rats, both the reproductive capacity and the frequency of regular estrous cycles are decreased compared to their young adult counterparts **(1)**. The age-associated changes in estrous cyclicity observed in M-A rats are associated to a delay as well as an attenuation of the preovulatory Luteinizing hormone (LH) surge **(2)**, increased LH serum follicle-stimulating hormone (FSH) levels, and altered circulating estradiol profiles with absence of the surge in the noon of the proestrus with respect to the young rats, **(3)**. There is clear evidence that in these animals, ovarian failure is preceded by the appearance of a diminished response of the neuroendocrine axis to positive feedback from estradiol **(4,5)**. Our group has implemented strategies capable of prolonging reproductive function in female rats. In female MA rats we demonstrated that intrahypothalamic gene therapy for insulin-like growth factor-I (IGF-I) started at 6 months of age extends the regular cyclicity of the animals beyond 10 month (the age at which MA rats stop ovulating) and preserves the integrity of the ovarian structure. At 11 months of age, the treated rats showed, in general, a cyclicity preserved in its regularity as well as a normal ovarian histology, while the controls were, at the same age, mostly acyclic and exhibited a high percentage of polycystic ovaries and few corpora lutea **(6)**.

### Aging and epigenetics

The discovery of animal cloning **[7,8]** and the subsequent development of cell reprogramming technology by means of the four reprogramming factors, Oct4, Sox2, Klf4, c-Myc, also known as the Yamanaka genes, **[9]** ushered in a technological and conceptual revolution that led to the achievement of cell rejuvenation by full reprogramming and the emerging view of aging as a reversible epigenetic process where cumulative DNA damage does not appear to play a central role as was thought for a long time **(10)**.

#### Regenerative effects of Yamanaka gene therapy

It is of significant interest that Yamanaka gene therapy in the retina of an experimental-glaucoma model and in middle-aged mice ameliorated their visual acuity**[13]**.. In progeric mice, transgenic for OSKM factors, cyclic partial reprogramming attenuated several signs of aging in visceral tissues and extended by 50% the survival of experimental versus control counterparts **[11]**. Furthermore, in a very recent preprint **[12]** it was shown that intravenous gene therapy with a regulatable AAV9 vector system that expresses the OSK genes, implemented in senile mice (29.2 months), prolonged the survival of the animals by two months compared to control counterparts. This is the first report showing that Yamanaka gene therapy in WT mice can extend their survival. We have recently observed that 39-day OSKM gene therapy in the dorsal hippocampus of old rats significantly reversed the typical learning deficits displayed by aged rats **[unpublished results]**.

This line of evidence prompted us to implement regenerative gene therapy in the hypothalamus of young female rats using an adenovector constructed by us that expresses a polycistronic cassette (the STEMCCA system) harboring the OSKM genes as well and GFP gene, all under the control of a bidirectional regulatable promoter **(14)**. We hypothesized that the long-term expression of the OSKM genes in the hypothalamus of young female rats could slow down the rate of decline of fertility in the animals as they near the age of ovulatory cessation (10 months in our rat colony). We report here that 9 months old female rats submitted to hypothalamic OSKM-GFP gene therapy at 4 months of age, display a higher fertility than control vector-treated counterparts.

## Materials and Methods

### Adenoviral Vectors

#### Construction of a regulatable HD-recombinant adenoviral vector (RAd) Tet-Off adenovector harboring the GFP and Yamanaka genes

The adenovector was constructed using a commercial kit (Microbix Inc., Ontario, Canada) that provides the shuttle plasmid pC4HSU, the helper adenovirus H14 and the HEK293 Cre4 cell line. A full description of the adenovector constructed has been already documented **(10)**. Briefly, we cloned a construct harboring the bicistronic tandem Oct4-f2A-Klf4-ires-Sox2-p2A-cMyc (known as hSTEMCCA, generously provided by Dr. G. Mostoslavsky, Boston University) under the control of the bidirectional promoter PminCMV-TRE-PminCMV a regulatable Tet-Off promoter which on its second end is flanked by the gene for humanized green fluorescent protein (hGFP).

The hSTEMCCA cassette harbors the 4 Yamanaka genes grouped in pairs placed downstream and upstream of an internal ribosome entry site (IRES). In turn, each pair of genes is separated by a type 2A CHYSEL (cis-acting hydrolase element) self-processing short sequence which causes the ribosome to skip the Gly-Pro bond at the C-terminal end of the 2A sequence thus releasing the peptide upstream the 2A element but continuing with the translation of the downstream mRNA sequence. This allows near stoichiometric co-expression of the two cistrons flanking a 2A-type sequence **[15]**. The whole expression cassette (STEMCCA cassette 10,065 bp) was cloned into the pC4HSU HD shuttle at the AscI and SwaI sites giving rise to pC4HSU-STEMCCA-tTA **(Fig 1)**. The pC4HSU HD shuttle consists of the inverted terminal repeats (ITRs) for Ad 5 virus, the packaging signal and part of the E4 adenoviral region plus a stuffer non-coding DNA of human origin which keeps a suitable size (28-31 Kbp) of the viral DNA.

**Figure 1.**
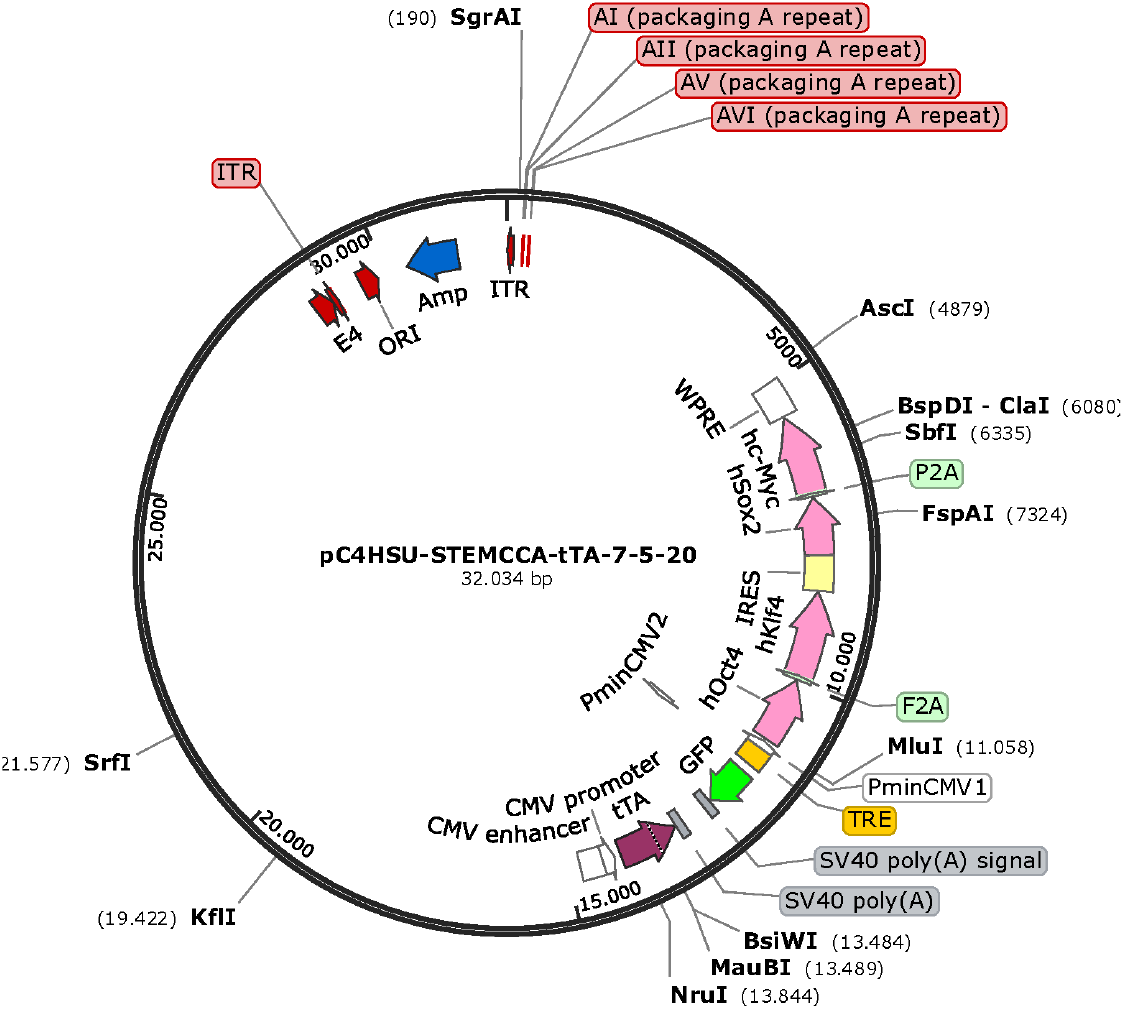
pC4HSU HD shuttle-. It consists of the inverted terminal repeats (ITRs) for Ad 5 virus, the packaging signal and part of the E4 adenoviral region plus a stuffer non-coding DNA of human origin which keeps a suitable size (28-31 Kbp) of the viral DNA. The whole expression cassette (STEMCCA cassette 10,065 bp) was cloned into the pC4HSU HD shuttle at the AscI and SwaI. For further details see **Fig 2 A** legend.

The linearized DNA backbone of the new HD-RAd **(Fig. 2A)** was transfected in Cre 293 cells. For expansion, the helper H14 adenovirus was added to the cell cultures at a multiplicity of infection (MOI) of 5. In H14, the packaging signal is flanked by lox P sites recognized by the Cre recombinase expressed by the 293 Cre4 cells. Therefore, the helper virus provides in trans all of the viral products necessary for generation of the desired HD-RAd. The infected 293 Cre4 cells were left for 2-3 days until cytopathic effect (CPE) was evident. Cells and medium were collected and submitted to 3 freeze-thaw cycles to lyse them. Clear lysates were obtained, mixed with H14 helper virus and added to a fresh culture of Cre4 293 cells at MOI 1. When CPE appeared, passage 2 (P2) cell lysates were prepared. This iterative co-infection process was carried on five more times in order to generate enough HD-RAd particles as to proceed to the purification step. The newly generated HD-RAd was rescued from P5. The HD-RAd so generated was purified by ultracentrifugation in CsCl gradients. Final virus stock was titrated by lysing the viral particles, extracting their DNA and determining its concentration in a Nanodrop spectrophotometer. For the first preparation, titer was 12 × 10^11^ physical viral particles (pvp) /ml.

**Figure 2.**
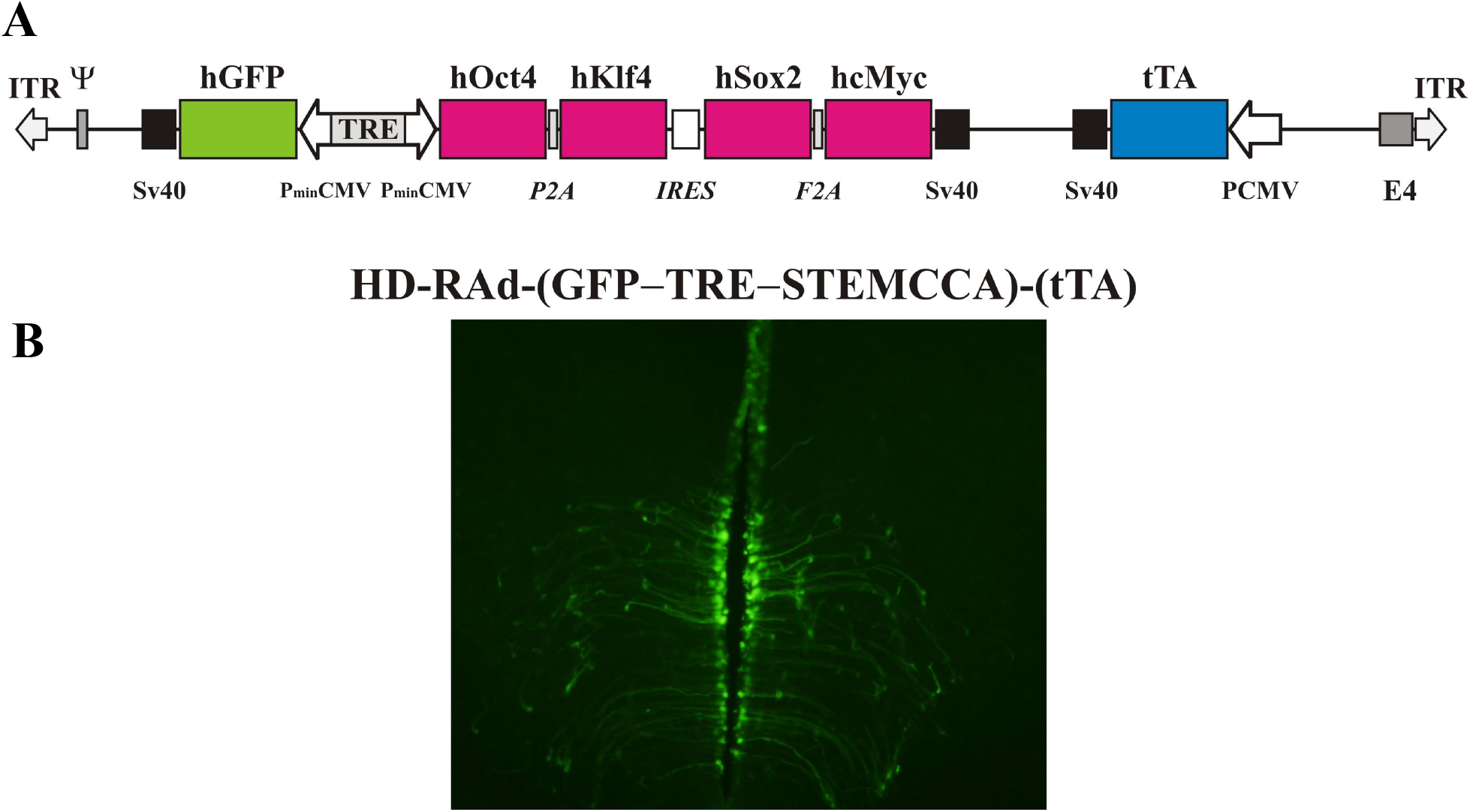
**Panel A** The figure illustrates the basic components of HD-RAd-STEMCCA-GFP-Tet-Off genome. Abbreviations- GFP: humanized Green Fluorescent Protein; TRE: Tetracycline responsive element; tTA: chimeric regulatory protein; PminCMV: cytomegalovirus minimal promoter; SV40pA: polyadenylation signal; ITR: inverted terminal repeats; Ψ: packaging signal. **Panel B-** Expression of GFP from the adenovector HD-RAd-STEMCCA-GFP in the third ventricle of the rat hypothalamus 6 days after vector injection in the lateral ventricles. Notice the processes of the ependymal tanycytes forming an extensive network in the hypothalamic parenchyma. Obj. 10X

We also constructed two control regulatable HD-RAd Tet-Off harboring either the hGFP gene or the gene for the DsRed2, red fluorescent protein isolated from Discosoma sp.

### Animals and in vivo procedures

Female Sprague-Dawley rats aged 3-4, 10 and 26 months were used. The animals were raised in our institution (INIBIOLP) and housed in a temperature-controlled room (22 ± 2ºC) on a 12:12 h light/dark cycle (lights on from 7 to 19 o’clock). Food and water were available ad libitum. All experiments with animals were performed according to the Animal Welfare Guidelines of NIH (INIBIOLP’s Animal Welfare Assurance No A5647-01).

#### Stereotaxic injections

Rats were anesthetized with ketamine hydrochloride (40 mg/kg; ip) plus xylazine (8 mg/kg; im) and placed on a stereotaxic apparatus. In order to access the,mediobasal hypothalamus (MBH) the tip of a 26G needle fitted to a 10μl syringe was brought to the following coordinates relative to the bregma: 3.0 mm posterior, 8.0 mm ventral and 0.6 mm right and left **(16)**.

#### Vaginal smears

Vaginal secretion was collected daily, between 11 and 13 o’clock, with a plastic pipette filled with 20 μl normal saline (NaCl 0.9%) by inserting the tip into the rat vagina, but not deeply. A drop of vaginal fluid was smeared on a glass slide and the unstained material was observed under a light microscope, with a 40X phase-contrast objective. Three types of cells can be recognized: round and nucleated ones are epithelial cells; irregular ones without nucleus are cornified cells; and little round ones are leukocytes. The proportion among them was used for determination of the estrous cycle phases **(17, 18)**, which are indicated as follows, P, proestrus; E, estrus, M, metestrus; D, diestrus; Pe, proestrus entering estrus; Dp, Diestrus entering proestrus.

In M-A rats spending several days in a row in constant estrus (CE), a CE cycle was defined, for quantitation purposes, as a period of 5 consecutive days of vaginal smears showing only cornified cells. For instance, if an animal spent 13 days in a row in CE they were counted as 13/5=2.6 CE cycles.

### Assessment of the effect of age on cycle regularity

Females aged 3, 10 and 26 months were submitted to daily vaginal smears sampling for 30 days.

#### Experimental design for long-term OSKM-GFP gene therapy in young females

Four-month old cycling females were allotted to a control or experimental group of 12 animals each, thus forming 2 groups: Control, HD-RAd-GFP-injected control (**GFP group**) and HD-RAd-OSKM-GFP-injected experimental (**OSKM group**). Occasionally, 7-day vaginal smears were collected to verify that GFP and OSKM rats were cycling regularly (data not shown).

When the GFP and OSKM rats reached the age of 9 months and 10 days (40 weeks), a group of 12 intact female rats (3-4 mo.) was added (**Y group**). In each experimental group the 12 animals were divided into 4-rat subgroups, each of which was mated for one week with a young male rat (3-4 mo). At the end of the pregnancy period pregnant rats were placed in single cages the number of pups per litter was counted and the newborn rats were weighed.

### Hypothalamic section assessment by fluorescence microscopy

Rats were placed under deep anesthesia and perfused with phosphate buffered formaldehyde 4%, (pH 7.4) fixative. Each brain was removed and trimmed down to a block containing the whole hypothalamus. The block was then serially cut into coronal sections 40 μm thick on a vibratome (Leica, Nussloch, Germany).

Hypothalamic brain sections were mounted with Fluoromount water-soluble mounting medium. Images of hypothalamic and other brain sections were captured using an Olympus DP70 digital camera attached to an Olympus BX51 fluorescence microscope (Tokyo, Japan). Digital images were analyzed using the ImagePro Plus (IPP™) v5.1 image analysis software (Media Cybernetics, Silver Spring, MA).

### Statistical Analysis

The t-test or analysis of variance (ANOVA) was used, as appropriate, to evaluate group differences. Tukey’s method was chosen as a post hoc test.

## RESULTS

### HD-RAd OSKM-GFP expression in the hypothalamus of young rats

Both, the HD-RAd OSKM-GFP experimental vector and the HD-RAd Tet-Off-control vectors were stereotaxically injected in the MBH of young female rats. Six days after injection, hGFP expression of the OSKM-GFP vector was strong. The processes of the ependymal tanycytes formed an extensive network in the hypothalamic parenchyma (**Fig 2-Panel B**). When our control vector HD-RAd-DsRed2 vector was injected into the hypothalamic parenchyma of young rats, DsRed 2 expression remained high for at least 30 days **(Fig 3)**.

**Figure 3.**
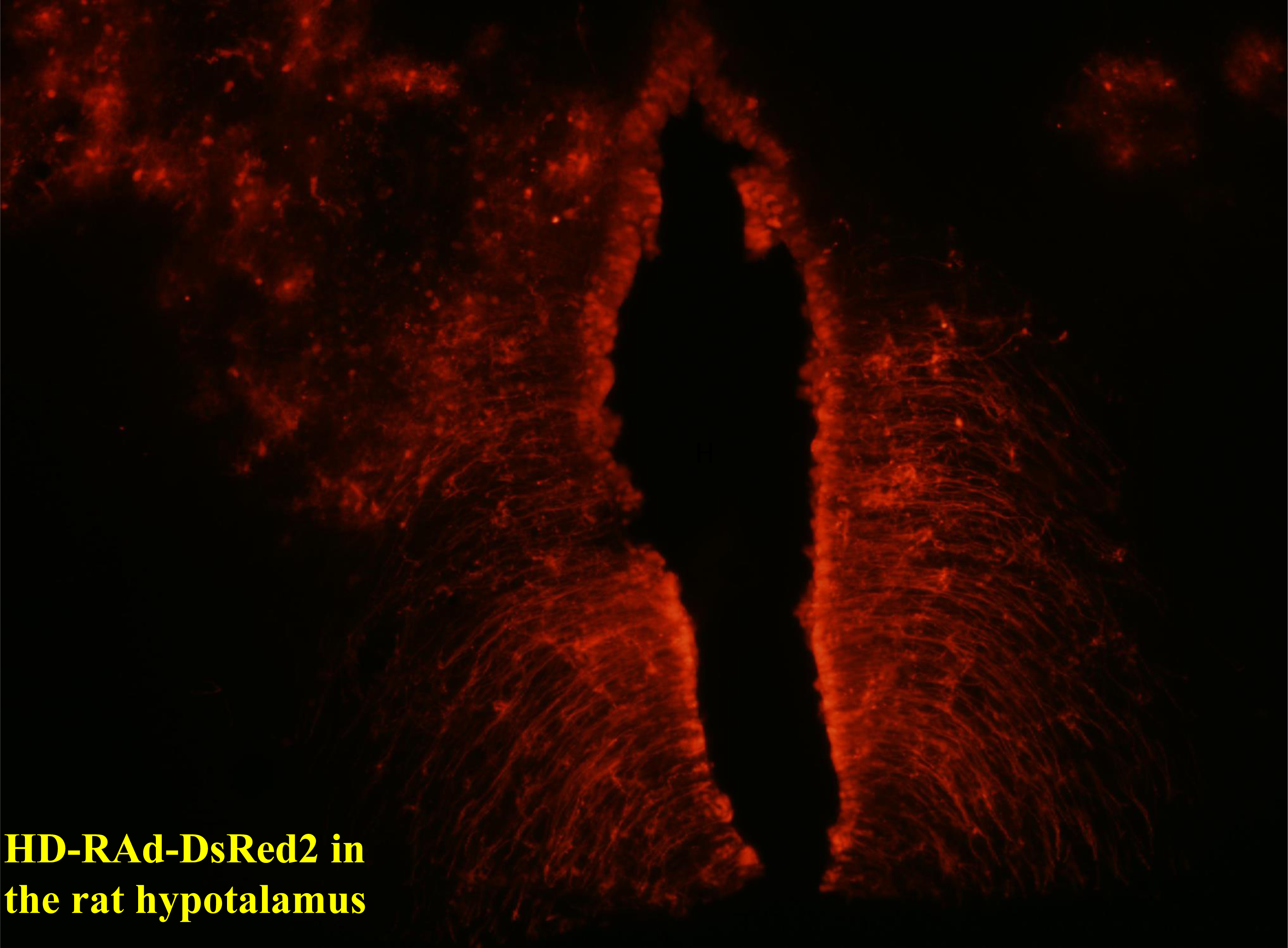
Expression of DsRed2 from the control adenovector HD-RAd-DsRed2 in the rat hypothalamus 30 days after vector injection. Notice the processes of the ependymal tanycytes forming an extensive network in the hypothalamic parenchyma. Obj. 10X

### Characterization of estrous cycle patterns throughout the lifespan

Initial characterization in our female rat colony of estrous cycle patterns from youth through very old age showed that transition from regular to irregular cyclicity takes place at around 9 months of age and is followed by a prevalence of CE status from age 10 to 18-20 months **(Fig. 4)**. Animals older than 20 months progressively go into a constant diestrus phase. Thus, 3 month old females typically show 4-5-day estrous cycles characterized by one day or less of P (in some cycles P was so short that when vaginal smear was performed the cell proportion was consistent with proestrus entering estrus or sometimes just estrus), one day of E and 2-3 days of M and/or D. Ten-month old females typically show a prevalence of lengthy CE periods with interspersed irregular cycles. Their ovarian histology shows numerous follicular cysts and scarce CL (data not shown). In senescent females (26 mo.) CE gave way to an increasing incidence of long periods of D with occasional 2-3 day-periods of P and E vaginal cytology as well as some CE periods. Senescent ovaries showed follicular cysts and some CL (data not shown).

**Figure 4.**
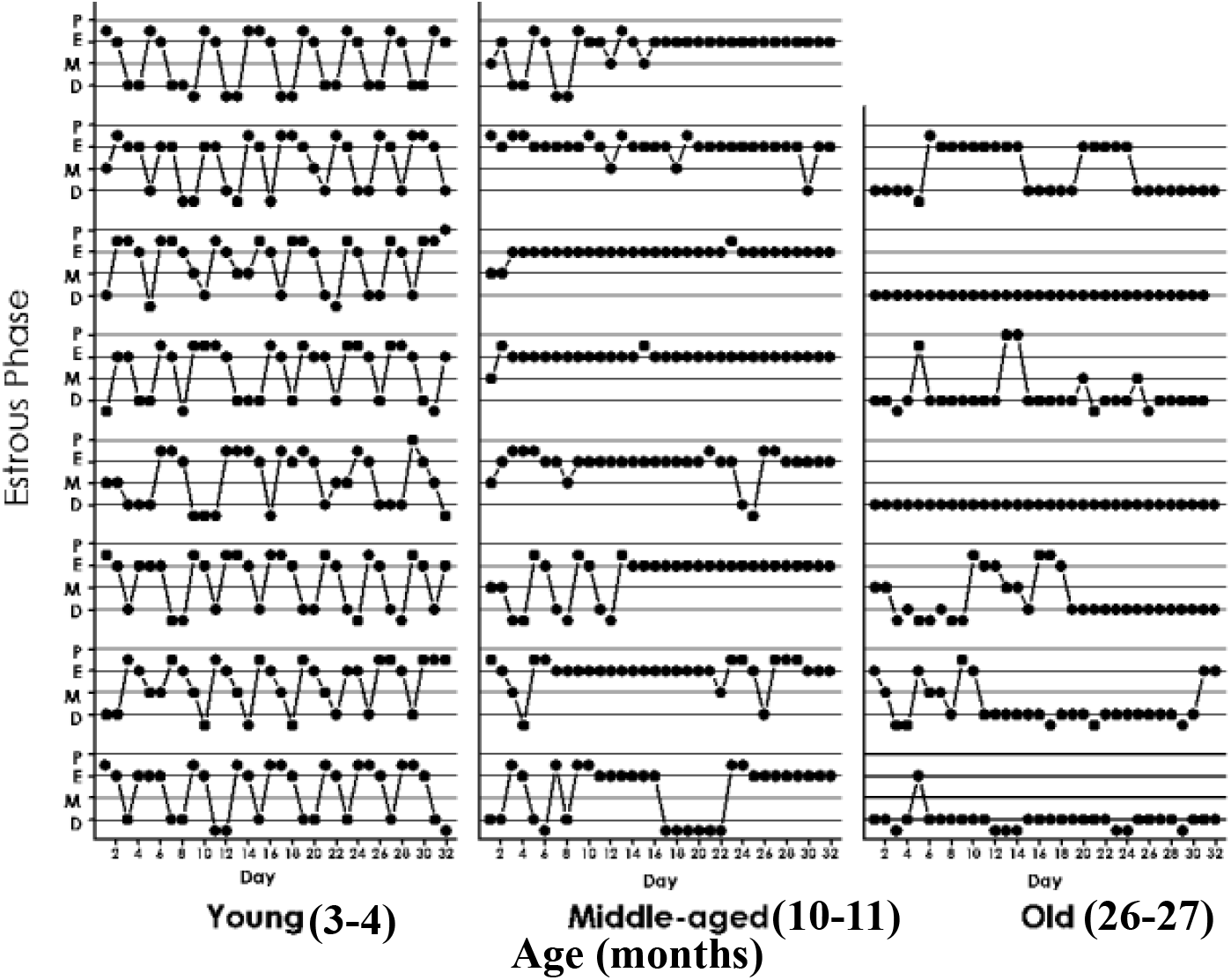
Estrous cycle patterns in young (3 mo., A), middle-aged (10 mo., B) and old (26 mo., C) female rats. The Y axis represents the estrous cycle phases of the animal on the indicated day of a 31-day window (X-axis) as determined by vaginal cytology. Four estral phases were defined according to the proportion of epithelial and cornified cells or leukocytes; Proestrus (P) was characterized by abundance of epithelial cells and the presence of leukocytes. Estrus (E), showed abundance of cornified cells and to a lesser extent epithelial cells. Metestrus (M), was defined by the prescence of an increasing number of leukocytes accompanied by substantial numbers of cornified cells. Diestrus (D) was characterized by high abundance of leukocyes which constituted the vast majority of cells present in the vaginal smears at D. Two transitional stages were also identified, Pe, charecterized by abundance of epithelial cells, almost no leukocytes and some cornified cells; Dp, characterized by abundance of leukocytes and some epithelial cells.

### Effect of hypothalamic OSKM gene therapy on fertility in M-A female rats

As expected, the young group showed a much higher pregnancy rate (83%) than the M-A controls (8.3%). Interestingly, M-A OSKM treated rats showed a 25% rate of fertility, which although far lower than in young rats, it was 3 times higher than in M-A controls. **(Fig. 5, upper plot)**. Also as expected, the litter size was higher in the young (mean litter size 9.1 pups) than in the M-A controls **(Fig. 5, middle plot)** and OSKM-treated (mean litter size, 3 pups in each group). The body weight of the pups was lower in the young rats (mean BW in pups from young rats, 9.2 g) than in the M-A controls and treated (mean BW, 11.6 and 11 g, respectively). **(Fig. 5, lower plot)**. The total number of pups born from Y, MA-c and MA-oskm was 83, 3 and 8, respectively. All pups survived and showed similarly normal behavior in the three groups.

**Figure 5.**
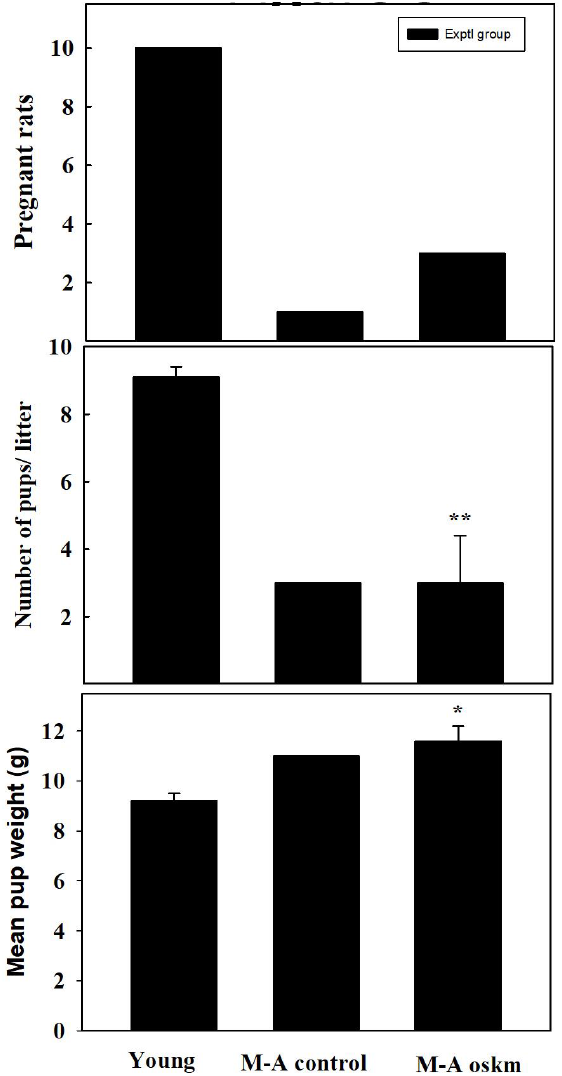
Effect of long-term OSKM-GFP gene therapy on the pregnancy rate, litter size and pup body weight on M-A female rats. There are three experimental groups; 4 months old intact females **(Young)**, 9.3 months old control rats **(M-A control)** and 9.3 months old OSKM-treated rats **(M-A oskm)**. Pups were weighed when they were 1 or 2 days old.

## DISCUSSION

Although the sequence of changes that take place during reproductive aging in female rats is qualitatively similar in most strains, the timing is likely to differ among strains and in different laboratory environments. This is why we considered it necessary to characterize the chronology of reproductive changes in our female rat colony before attempting to implement long-term protective gene therapy in young animals. In qualitative terms, the age changes in vaginal cytology observed in our Sprague-Dawley females are in agreement with early reports in Long-Evans rats. Thus, in Long-Evans females the vaginal smears show regular 4-5 day cycles from 2 to 10 months of age, transitioning to irregular cycles and persistent vaginal cornification (CE) during the following two months **(19, 20)**. In our M-A females this transition takes place earlier. After 18-20 months, most of the animals progressively become pseudopregnant (PSP), a condition characterized by persistent diestrus vaginal cytology interrupted at irregular intervals, by 2-3 days of nucleated and cornified vaginal smears. In Long-Evans females the PSP stage has been reported to occur between 27-30 months of age **(20)**. While the CE stage is associated with an anovulatory condition, PSP females show intermittent ovulatory activity **(21)**.

The M-A females were mated 5.8 months (174 days) post adenovector injection, which, despite the long-term expression of HD-RAds, makes it likely that OSKM gene expression in the MBH at the time of mating was significantly weaker than at the time of injection. We have previously demonstrated that in our female rat colony, ovulation remains regular until 9 months of age, becoming irregular at age 10 months **(6)**. Clearly, regular ovulation is a necessary but not sufficient condition for keeping the rats at an optimal fertility levels, progressive decline in other components of the reproductive system has a marked overall impact on fertility, as shown in the M-A controls. Considering that at the time of mating, M-A rats were nearing reproductive cessation, the 25% fertility rate observed in the OSKM-treated rats represents a significant beneficial effect of the OSKM treatment on the rate of fertility. It may be possible that if rats had been mated at an earlier age, 7 months, for instance, the fertility rate would have been higher than 25% but perhaps not three times higher than the fertility of control counterparts. Other parameters like litter size and mean pup BW seem less sensitive to the beneficial effects of OSKM treatment than fertility.

It should be pointed out that only a small proportion of hypothalamic cells are expected to have been transduced by our OSKM-GFP vector. However, since an extensive network of tanycytic processes perfuses the hypothalamic parenchyma **(Fig. 2 B and 3)** it is likely that the influence of the relatively small number of transduced cells was amplified by this network. Those few cells expressing the OSKM genes may have generated a neuroprotective environment in the hypothalamus, effectively slowing down the effect of age on reproductive function.

It was not possible to assess the impact of the OSKM treatment on the hypothalamus of the rats after delivery, as pregnant rats and nonpregnant animals would have shown pregnancy-related differences in their hypothalamic DNA methylation profile. A possible way to assess the impact of OSKM gene therapy on hypothalamic DNAm age would be to perform a separate experiment, skipping mating, and sacrificing the rats 9 months after vector injection.

Our results are in line with the evidence that viral vector-mediated delivery of the Yamanaka genes in the brain has strong regenerative effects without adverse side effects **(13)**. The particular significance or the present results is that, for the first time, they show that long-term OSKM gene therapy in the hypothalamus is able to extend the functionality of such a complex system as the hypothalamo-pituitary-ovarian axis.

## Acknowledgements

The authors thank Mr. Mario R. Ramos for design of the figures and Ms. Yolanda E. Sosa for technical and editorial assistance

## Funding

Our studies are supported in part by research grant Grant # SEGR-9-23-21 from the Society for Experimental Gerontological Research (SEGR) and grant PICT 2018-00907 to RGG

## Competing Interests

None of the authors has potential competing interests to disclose.

## Notes

### Competing Interest Statement

The authors have declared no competing interest.

